# Comprehensive Benchmarking and Integration of Tumour Microenvironment Cell Estimation Methods

**DOI:** 10.1101/437533

**Authors:** Alejandro Jiménez-Sánchez, Oliver Cast, Martin Miller

**Affiliations:** Cancer Research UK Cambridge Institute, University of Cambridge, Li Ka Shing Centre, Robinson Way, Cambridge CB2 0RE, UK

## Abstract

The tumour microenvironment comprises complex cellular compositions and interactions between cancer, immune, and stromal components which all play crucial roles in cancer. Various computational approaches have been developed during the last decade that estimate the relative abundance of different cell types in an unbiased manner using bulk tumour RNA data. However, a comparison that objectively evaluates the performance of these approaches against one another has not been conducted. Here we benchmarked six widely used tools and gene sets: Bindea *et al.* gene sets, Davoli *et al.* gene sets, CIBERSORT, MCP-counter, TIMER, and xCell. We also introduce Consensus^TME^, a consensus approach that uses the union of genes that the six tools used for cell estimation, and corrects for tumour-type specificity. We benchmarked the seven tools using TCGA DNA-derived purity scores (33 tumour-types), methylation-derived leukocyte scores (30 tumour-types), and H&E deep learning derived lymphocyte counts (13 tumour types), and individual benchmark data sets (PBMCs and 2 tumour types). Although none of the seven tools outperformed others in every benchmark, Consensus^TME^ ranked consistently well in all cancer-related benchmarks making it the top performing method overall. Computational methods that provide robust and accurate estimates of non-cancerous cell populations in the tumour microenvironment from tumour bulk expression data are important tools that can advance our understanding of tumour, immune, and stroma interactions, with potential clinical application if high accuracy estimates are achieved.

## INTRODUCTION

The tumour microenvironment (TME) plays an active role in in tumour initiation, progression, metastasis, and treatment response. Thus, studying the TME is a central paradigm of cancer research. However, a great variety of stromal and immune cell types populate tumour tissues, and the complex interactions between these different components of the tumour microenvironment is still unclear. Traditionally, cells from the TME have been quantified using immunohistochemistry (IHC), immunofluorescence (IF), and flow cytometry, and more recently using cytometry by time of flight (CyTOF). These methods, although accurate, are laborious, low throughput and require pre-selected cellular markers, making their application in large number of samples and measurements challenging. Thus, their systematic application for comprehensively investigating the various different cell types in the TME in an unbiased manner is limited. Single cell RNA sequencing (scRNA-seq) has begun to fill this gap, however scRNA-seq is still too expensive to apply on large number of samples such as The Cancer Genome Atlas (TCGA) which consists of thousands of genomically profiled tumour samples which are also clinically well annotated. The study of the different cell subpopulations of the TME in TCGA has become an important goal, but also an important challenge for bioinformatics, since cell-type information identity is mixed in bulk tumour transcriptomics data.

Estimation of non-cancerous cell proportions from bulk tumour samples has been performed using genomics data such as whole-exome sequencing, microarrays, RNA-seq, or DNA methylation data. During the last decade, multiple computational approaches have been developed intending to quantitatively or semi-quantitatively calculate distinct TME cell-type population estimates^1^. A variety of statistical frameworks and algorithmic procedures have been employed, and each method has used different benchmark datasets^1^. In general, two different algorithmic classes into which most methods can be classified are: deconvolution algorithms and gene set enrichment-based methods. Importantly, both classes rely on cell-type specific markers that are selected according to prior knowledge. The deconvolution algorithms use linear combinations of the expression values of the cell-specific genes, while gene set enrichment-based methods rank the genes of a mixture sample and compute enrichment scores as a function of the ranked selected genes. This is a challenging task, particularly to reliably estimate lowly abundant cell populations, and also because the gene sets are not unique to any particular cell type. Thus, there is not a straightforward solution for accurate TME cell estimation, and one of the problems in the field is that each method has claimed to outperform others in their own benchmarking experiments^2,3^. Thus, the need for independent and more comprehensive benchmarks has been pointed out^4^. Here, we developed a consensus approach (Consensus^TME^) that leverages current state-of-the-art knowledge by compiling common cell-type specific gene sets used by six published TME cell estimation methods. We performed pan-cancer benchmarks using publicly available bulk RNA sequencing data from TCGA and side-by-side comparisons using the methods’ independent benchmarks. The Consensus^TME^ approach is evolvable by design allowing new methods and algorithms to be incorporated and their performance compared with continuously updated benchmark data sets. Overall, the consensus approach will lead towards robust and improved tools for the estimation of cell-type quantities using bulk expression data of human tumour samples.

## RESULTS

### Consensus tumour-type specific superset of tumour microenvironment cell populations

Following the generation of large data sets of tumour genomic profiles such as TCGA and ICGC, various computational tools assessing TME cell populations have been developed, each using different algorithms, gene markers, and validation benchmarks. To build on the knowledge of cell-type specific gene sets represented in the diversity of these methods, we sought an integrative strategy that incorporates knowledge from existing tools. Consensus^TME^ integrates cell-type specific gene markers from independent cell estimation methods and uses single sample gene set enrichment analysis (ssGSEA) to compute TME cell-type and tumour specific enrichment scores from bulk expression data (Figure 1A). The ssGSEA approach was selected because its treats microarray and RNAseq values in the same way, since it is based on the ranked genes rather than the actual values. To generate Consensus^TME^ gene sets, we selected six widely used cell estimation methods: CIBERSORT^5^, TIMER^6^, MCP-counter^7^, xCell ^8^, and the gene sets generated and use in Bindea et al.^9^ and Davoli et al.^10^ here called “Bindea” and “Davoli” supersets, respectively. In brief, we first selected cells that are estimated by at least two methods. Second, we generated a gene set for each cell type by using the union of genes used by the methods to estimate that cell type, and removed genes that correlate (rho > −0.2) with tumour purity^6^ (see Methods). Therefore, Consensus^TME^ aggregates cell-type specific genes that have been independently considered relevant by different methods, and estimates their abundance in a tumour type specific manner.

**Figure 1:**
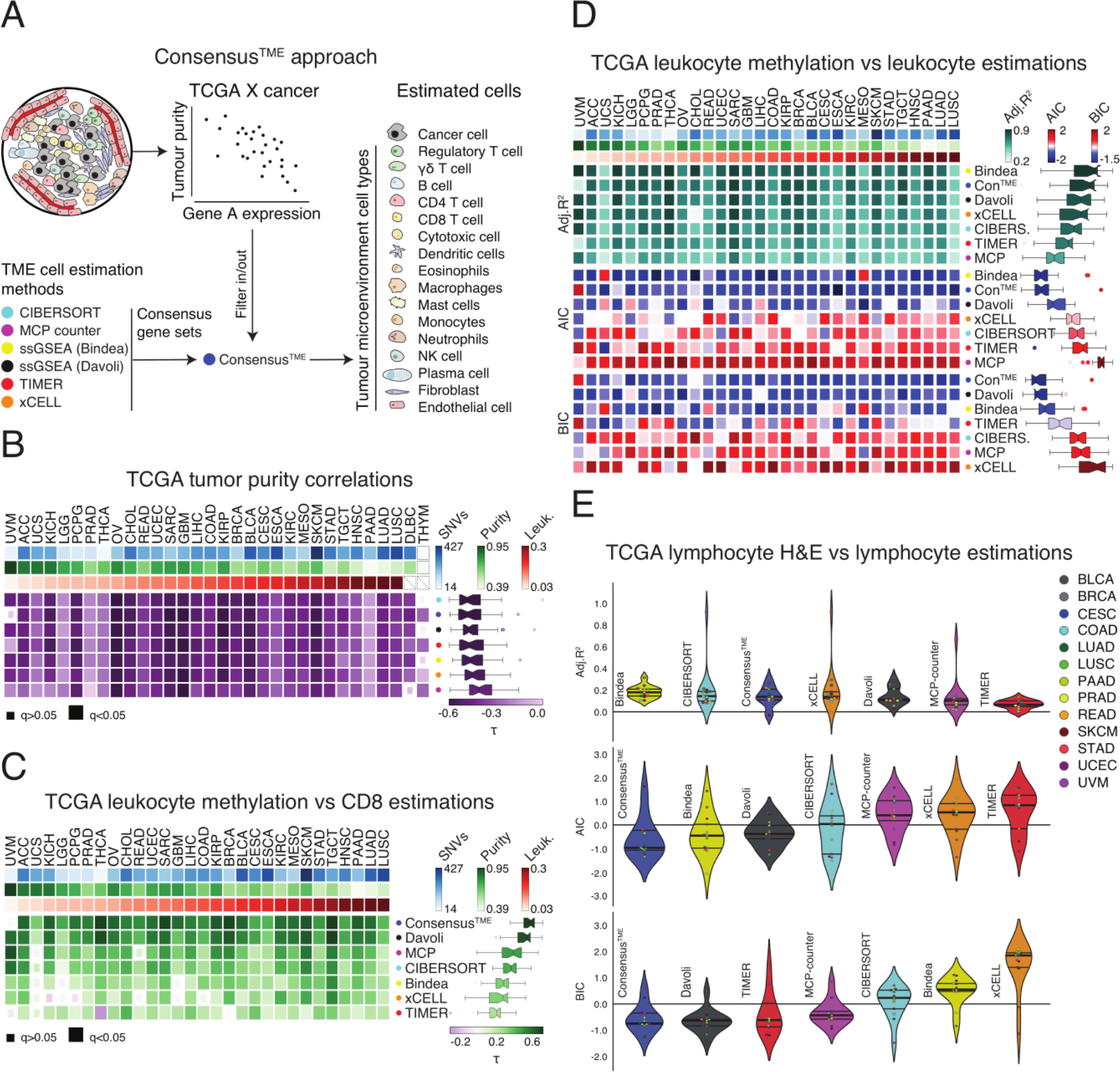
Benchmark of methods for estimating TME cell components using purity and leukocyte data from TCGA data. A) Bioinformatic tools benchmarked, and Consensus^TME^ development strategy (see Methods). B) Kendall’s correlation coefficients (τ) of DNA-derived ABSOLUTE purity scores^11,12^ and RNA-derived estimated purity by the different methods. ESTIMATE’s stromal scores were added to all the methods to account for stromal quantities, since not all methods analyse stromal cells. C) Kendall’s correlation coefficients (τ) of methylation-derived leukocytes’ scores^12^ and RNA-derived CD8^+^ T cell estimations. D) Multiple linear regression models of leukocyte methylation scores as response variable^12^ and RNA-derived leukocyte estimations as explanatory variables. Column heatmaps are sorted according to leukocyte methylation scores (Left: Low, Right: High), rows are sorted according to median performance (Top to Bottom: Decreasing performance). E) Multiple linear regression models of deep learning H&E-derived lymphocyte counts as response variable^14^ and RNA-derived lymphocyte estimations as explanatory variables. Kendall’s correlation significance is shown with small/large squares as indicated by the q-value multiple test corrected. Adjusted R^2^, Akaike Information Criterion (AIC) z-scores, and Bayesian Information Criterion (BIC) z-scores were compared across models generated by each tool cellular estimations. Lower AIC and BIC values represent a better goodness-of-fit penalising the number of variables. Median single nucleotide variants (SNVs), ABSOLUTE purity scores (purity), and leukocyte methylation scores (Leuk.) per tumour type are shown in B, C, and D. Violin plots are sorted according to median correlation coefficient (Left to Right: Decreasing performance).

### Pan-cancer leukocyte and lymphocyte benchmarks

To benchmark the different methods in an objective and systematic manner, we used publicly available data from between 13 and 32 tumour types comprising 9,142 tumour samples in total. First, to evaluate the ability of each method to capture the overall amount of immune component in the TME we correlated DNA-based tumour purity scores^11,12^ with immune scores inferred by the different methods either natively, or derived when not generated by default (see Methods). Since tumour purity does not account only for immune cell infiltration but also for other stromal cells (e.g. fibroblasts and endothelial cells), this would affect the correlation of methods that only estimate immune cells. Therefore, we inferred stromal non-immune related content of all samples using ESTIMATE^13^ and added this value to all methods’ immune scores to create a purity score. We found that all six methods and Consensus^TME^ perform very similar to each other, with CIBERSORT, Consensus^TME^, and Davoli as the top 3 pan-cancer negative correlations (Figure 1B). Across tumour types, the different methods performed similar and very few correlations were not statistically significant. However, cancer-specific performance of methods was observed with methods consistently performing well in cancers such as ovarian cancer (OV) and glioblastoma multiforme (GBM) while showing lower performance in pancreatic adenocarcinoma (PAAD). Variation in performance was largely independent of cancer cellularity, mutation load, leukocyte fraction, and sample size for any of the methods.

We further evaluated the performance of the methods by using leukocyte fractions derived from methylation data for 30 tumour types^12^. For the initial analysis we assessed performance by correlating levels of CD8^+^ T cells, an important prognostic cell type, with leukocyte fraction. The best performing methods for this analysis were Consensus^TME^, Davoli and MCP-Counter. To extend this analysis to account for accuracy across multiple cell types we fitted multiple linear regression models using the leukocyte fraction as a response variable and only cell type estimates in the category of being leukocytes as explanatory variables for each method. Since different methods estimate different number of leukocytes, the coefficient of determination can be artificially increased by the number of variables in a model (i.e. overfitting). Thus, to more appropriately compare the models in an unbiased way we used adjusted coefficients of determination (R^2^), the Akaike information criterion (AIC), and Bayesian information criterion (BIC) for model comparison. These penalise model complexity (i.e. number of cell-types used in the models) to varying degrees. When comparing the R^2^ of the different models, the best performing methods were Bindea, Consensus^TME^, and Davoli (Figure 1D). Similarly, AIC and BIC scores showed that Consensus^TME^, Davoli, and Bindea models perform better than the other methods (models with lower AIC and BIC values are preferred). We also implemented multiple linear regression analysis using tumour-infiltrating lymphocyte counts derived from digitised H&E-stained images analysed through a deep-learning convolutional neural network approach^14^. Although low coefficient of determination values was obtained across methods, likely due to the very difficult task of computationally detect leukocytes on H&E images, the methods that obtained a higher coefficient of determination were Bindea, CIBERSORT, and Consensus^TME^, while the lowest AIC and BIC values were obtained by Consensus^TME^, Davoli, and TIMER (Figure 1E). Together, these broad pan-cancer benchmarks show a variation in the performance of the different methods when compared to each other, and no single method consistently outperforms the others. The cancer specific performance observed in these experiments was also an important finding that should be taken into account when considering the appropriateness of using these tools.

### Independent methods benchmarks

All methods tested, except Bindea and Davoli gene supersets, performed their own independent benchmarks in the original publications. Thus, we collected benchmarking data for CIBERSORT, xCELL, TIMER, and MCP-counter to carry out a side-by-side comparisons. We used the CIBERSORT benchmark data that consisted of peripheral blood mononuclear cells (PBMCs) of 27 human subjects quantified by flow cytometry^5^. Correlations between estimated immune cell types and the flow cytometry fractions showed that the best performing methods were MCP-counter, CIBERSORT, and xCELL (Figure 2A). However, most of the correlations lacked statistical significance, and due to the different cell types estimated by the different methods it is difficult to reach a conclusion. Similarly, the xCELL benchmark data set consisted of 16 PBMC leukocyte subsets from two different studies with 61 and 104 human subjects each, where PBMCs were measured using Cytometry by Time of Flight (CyTOF)^8^. Again, MCP-counter, CIBERSORT, and xCELL were the methods that showed best performance in these PBMC benchmarks, however many cell types did not reach statistical significance (Figure 2B).

**Figure 2:**
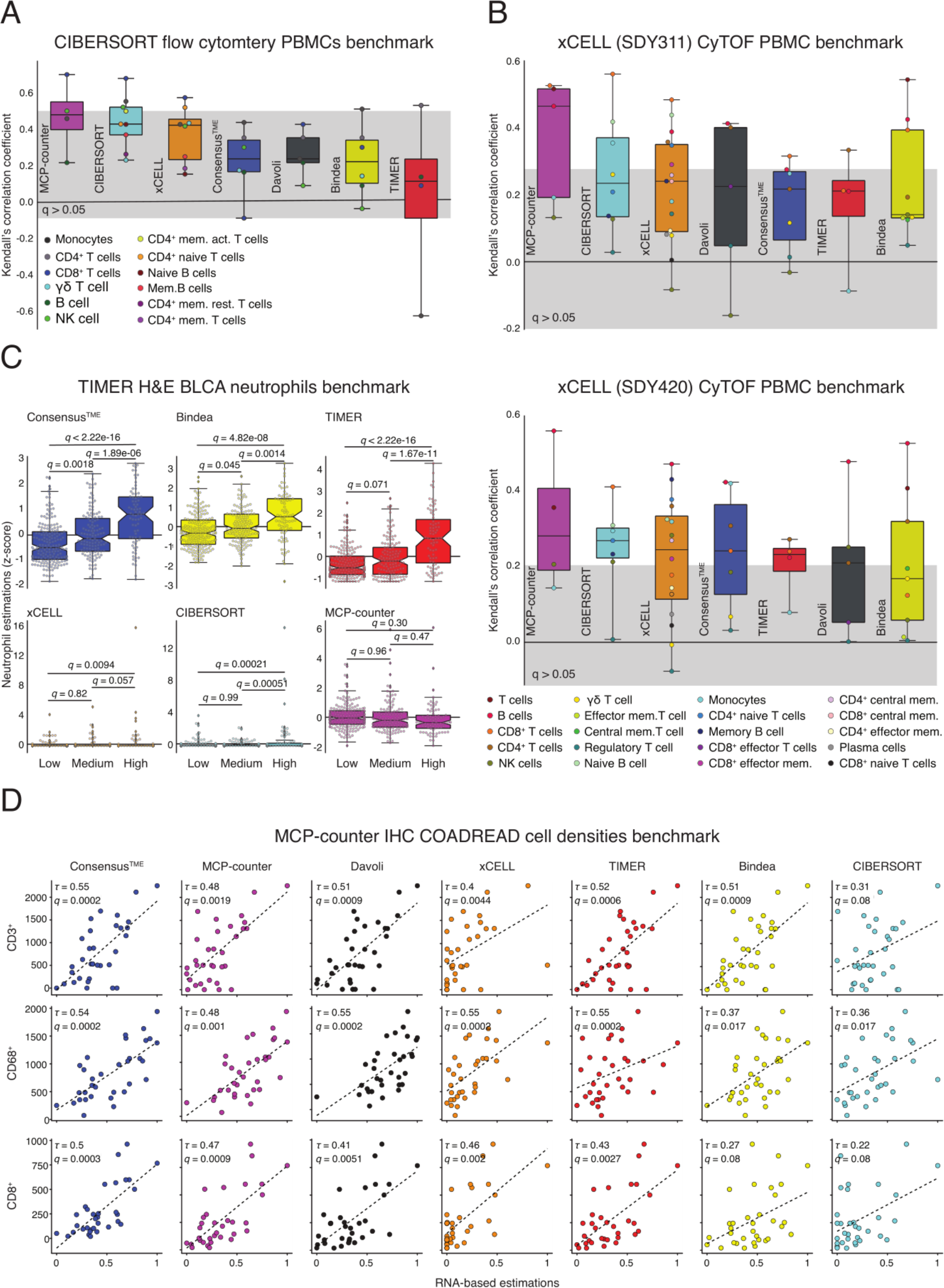
Side-by-side benchmark using original benchmarking datasets published by the individual methods. A) Kendall’s correlation coefficients (τ) of CIBERSORT PBMCs’ flow cytometry (n=20 samples) and B) xCell PBMCs’ CyTOF benchmarks. SDY311 (n=61 samples) and SDY420 (n=104 samples) are different benchmarks available from ImmPort^15^. Since not all methods estimate cell activation states, cell proportions were aggregated into the cell type and used for correlation with the methods that do not estimate the specific activation states. The grey box represents correlation coefficients that have a *q*-value > 0.05 (B-H method). Box plots are sorted according to median correlation coefficient (Left to Right: Decreasing Performance). C) Comparison between low, medium, and high categories of BLCA (n=404 samples) neutrophil H&E pathology counts. One-way ANOVA with Tukey HSD post hoc tests were employed to calculate *q*-values. Plots are sorted according to performance (Left-Top: Best, Right-Bottom: Worst). D) Kendall’s correlation coefficients (τ) of MCP-counter COADREAD IHC (n=38 samples) cell densities (cell/mm^2^). Plots are sorted according to median correlation coefficient (Left to Right: Decreasing Performance).

Finally, we used cancer-related benchmarks from TIMER^6^ and MCP-counter^7^. For TIMER’s benchmark, 404 TCGA bladder cancer samples were analysed by a pathologist who categorised them as low, medium, or high according to their neutrophil counts using H&E stained slides. Then performance was assessed by measuring the significance of difference between the computational estimates of samples in each category. Here, Consensus^TME^, Bindea, and TIMER obtained the best separation between categories, but only Consensus^TME^ and Bindea separated significantly the three categories after multiple test correction (Figure 2C). Interestingly, xCell, CIBERSORT, and MCP-counter were unable to differentiate the three categories, while Davoli does not estimate neutrophils. The MCP-counter benchmark consisted of IHC digital quantification of CD3^+^ (T cells), CD8^+^ (CD8^+^ T cells), and CD68^+^ (Monocytic lineage) cell densities from 38 colorectal cancer samples. Correlations between the methods’ estimations and the cellular fractions were computed (Figure 2D). Consensus^TME^, MCP-counter, and Davoli methods provided the best correlations, with Consensus^TME^ outperforming on the three cell types.

When observing the rank of methods across all benchmarking experiments (Supplementary figure S1) no one method was shown to consistently outperform all others. However, the integrative Consensus^TME^ approach was in the top three for all cancer-based benchmarks and achieved the lowest mean rank of all methods.

## DISCUSSION

With the recent generation of large publicly available molecular profiling of cancer samples, a variety of computational tools for analysis of cell components of the TME have been generated. In principle, the method of choice should be based on performance, however popularity and ease of use can also be reasons behind the method researchers select^4^. In the case of TME cell estimation from bulk expression data, this problem is magnified by the lack of objective and independent benchmark analyses, since most methods use their own benchmarks which may introduce biases and reliance on one type of data. Here we performed an unbiased and objective benchmarking exercise comparing six of the most widely used and recent tools, and also developed Consensus^TME^: a gene set enrichment-based method that integrates cells and genes from these six different tools in order to generate a consensus gene superset for each cell type that is tumour-type specific.

We performed pan-cancer benchmarks using orthogonal data types generated for TCGA samples. While DNA-derived tumour purity scores correlated negatively with RNA-derived TME estimations in all methods, leukocyte methylation scores showed some distinction across methods, and a lack of correlation in some cases. Also, different tumour types showed varying levels of correlation with leukocyte estimations, which could be due to the leukocyte methylation signature model. This was generated by comparing pure leukocyte cells and normal tissue methylation patterns with tumour-type specific methylation patterns^12^, and tumour infiltrating leukocytes may have different methylation patterns. Furthermore, lymphocyte deep learning H&E quantifications provided a lower association with lymphocyte RNA-derived estimations, an observation that has been reported before and considered to be in part due to the RNA-derived estimates reflect more cell counts, while spatial image-derived estimates reflect the fraction of lymphocytes per area^14,16^. Thus, this benchmark is inconclusive due to the uncertainty of both RNA-derived and imaged-based derived lymphocyte estimates, but it was included to achieve more comprehensive and orthogonal benchmarks. Moreover, similar results to the leukocyte methylation benchmark were detected. Side-by-side benchmarking on the different methods data sets showed that PBMC-based benchmarks present a low number of significant correlations across methods, and due to the diversity of cells tested and estimated by the different methods obtaining a concluding result out of these benchmarks is challenging. Moreover, for the application of TME cell estimation using bulk RNA tumour data, PBMC benchmarks may not be very informative as the transcriptomes of the circulating and tumour infiltrating immune cells are different. In contrast, both BLCA and COADREAD benchmarks on neutrophils, CD3^+^, CD68^+^, and CD8^+^ cells showed significant associations for some tools, particularly Consensus^TME^. These benchmarks showed that no independent method consistently outperforms other methods. Nevertheless, overall Consensus^TME^ ranked among the top three best performing methods in all cancer-relates benchmarks. Lastly, Consensus^TME^ is an evolvable method by conception, which means that other genes used by new methods can be added to existing supersets and tested with already established benchmarks, thus potentially improving its performance as new methods and gene sets are developed.

## METHODS

### Contact for Reagent and Resource Sharing

Further information and requests for resources should be directed to and will be fulfilled by the Lead Contact, Martin L. Miller (http://martin.miller@cruk.cam.ac.uk).

### Quantification and Statistical Analysis

#### Single-sample gene set enrichment analysis

Single-sample gene set enrichment analysis^17^, a modification of standard GSEA^18^, was performed on RNA measurements for each sample using the GSVA package version 1.28.0^19^ in R version 3.5.0 with parameters: method = ‘ssgsea’, and tau = 0.25. Normalized enrichment scores (NES) were generated for the hallmark gene sets^20^, immune and stromal signatures^13^, TME cell gene sets obtained from previous publications^9,10^, as well as the Consensus^TME^ gene sets (Figure S3A). Hallmark gene sets were obtained from MSigDB database version 6.1^20^.

#### Consensus^TME^

To generate the Consensus^TME^ gene sets we identified cell types that were deconvoluted by at least 2 different methods, 18 in total. We then combined the gene sets that the different methods considered for the deconvolution of such cell types. To include genes used in CIBERSORT, we first filtered out genes whose expression value was below 1.96 standard deviations of the mean for each cell type. In addition, we collapsed activated and resting states for corresponding cell types. The union of genes was the filtered to exclude genes whose expression has a Pearson’s correlation coefficient > −0.2 and a *p*-value > 0.05 when correlated with tumour purity; as defined by TIMER^6^. Finally, ssGSEA was employed to calculate NES for each cell type as described above. General immune scores for each tumour types were generated by combining the genes of the different immune cells into one gene set.

#### Comparison statistical metrics

Concordance between computational estimates and ground truth values was measured using either Kendall’s rank correlation coefficient or the multiple linear regression goodness of fit metrics: adjusted R-squared, Akaike information criterion (AIC), and Bayesian information criterion (BIC). AIC and BIC z-scores values were calculated to incorporate different tumour types in the comparisons since AIC and BIC values are unitless. Differences between groups of variables were identified using one-way ANOVA with Tukey honest significant differences post-hoc tests. All statistical tests were adjusted for multiple testing using the Benjamini-Hochberg procedure to control for false discovery rate (FDR).

#### TCGA immune estimations

RNA-sequencing (RNA-seq) data was collected from cBioPortal^21^. Batch normalisation had been applied and gene expression values calculated using the “RSEM” pipeline^22^. Four existing TME cell estimation methods and two published gene sets were used alongside Consensus^TME^ to produce relative abundances of immune cell types per sample across 32 tumour types. For each method, a general immune score was also derived if it was not already provided, representing the total level of immune cell infiltration in each tumour sample.

#### TME cell deconvolution methods

Cell deconvolution methods were used to estimate levels of non-cancerous cells in the TME. The methods employed were CIBERSORT^5^, MCP-counter^7^, TIMER^6^, xCell^8^, as well as gene sets collected from two previous publications^9,10^.

#### Bindea et al. and Davoli et al. gene sets

Gene sets provided by Bindea et al. and Davoli et al. were used with ssGSEA to provide enrichment scores for each of the immune signatures^9,10^. To generate general immune scores, genes selected for immune cells where combined into one gene set for each method independently.

#### xCell

The “xCell” R package (version 1.12) was used to generate immune estimates for the xCell method^8^. A general immune estimation score is already provided by xCell.

#### MCP-counter

Estimations for the MCP-counter method were produced using the “MCPcounter” R package (version 1.1.0)^7^. Immune scores for this method were produced in a similar manner as the ssGSEA methods by creating a union of signature genes for each of the cell types. The “MCPcounter.estimate” function was used to allow for the new signature.

#### CIBERSORT

CIBERSORT estimations were produced using the R source code, provided on request from the web resource^5^. CIBERSORT was run in “Absolute mode” (under beta development) using 100 permutations and quantile normalisation disabled as recommended for RNA-seq data. Absolute scores representing the “overall immune content” is produced natively by the algorithm in absolute mode.

#### TIMER

TIMER estimations were produced using R source code, available from the web resource^6^. Immune scores for TIMER were produced as a sum of the coefficients for each cell type.

#### Purity score benchmark

Pan-cancer purity scores were downloaded from the NIH Genomic Data Commons (https://gdc.cancer.gov/about-data/publications/PanCan-CellOfOrigin)^12^. Purity scores were generated using ABSOLUTE^11^ which uses copy number, variant allele frequency, and tumour specific karyotype data to calculate the cancer fraction of a tumour samples. To benchmark the immune estimation methodologies using purity of samples the immune scores were added to an independent stromal score; calculated through the use of ESTIMATE (version 1.0.13)^13^. ABSOLUTE’s derived tumour purity and the different methods tumour purity scores were correlated independently for each tumour type. ACC n=77, BLCA n=397, BRCA n=1052, CESC n=293, CHOL n=36, COAD n=281, DLBC n=47, ESCA n=162, GBM n=154, HNSC n=509, KICH n=66, KIRC n=498, KIRP n=285, LGG n=519, LIHC n=359, LUAD n=504, LUSC n=493, MESO n=81, OV n=293, PAAD n=159, PCPG n=165, PRAD n=473, READ n=92, SARC n=246, SKCM n=460, STAD n=403, TGCT n=155, THCA n=469, THYM n=103, UCEC n=175, UCS n=56, UVM n=80

#### Leukocyte methylation benchmark

Leukocyte methylation scores were downloaded from the NIH Genomic Data Commons (https://gdc.cancer.gov/about-data/publications/PanCan-CellOfOrigin)^12^. Leukocyte methylation and CD8^+^ T cell estimations were correlated, since CD8^+^ T cells are estimated by all the methods. Multiple linear regression was then employed using all leukocytes the methods estimate as explanatory variables and the leukocyte methylation scores as response variable. Leukocyte methylation data was log transformed to meet the normality and heteroscedasticity assumptions of the model. Adjusted R^2^, AIC, and BIC metrics were calculated to compare the goodness of fit between the methods while taking into consideration the number of variables included in the model. ACC n=79, BLCA n=405, BRCA n=1077, CESC n=306, CHOL n=36, COAD n=278, ESCA n=185, GBM n=137, HNSC n=522, KICH n=66, KIRC n=528, KIRP n=291, LGG n=526, LIHC n=372, LUAD n=505, LUSC n=499, MESO n=87, OV n=306, PAAD n=179, PCPG n=184, PRAD n=494, READ n=94, SARC n=263, SKCM n=471, STAD n=415, TGCT n=137, THCA n=509, UCEC n=177, UCS n=57, UVM n=80.

#### Somatic single nucleotide mutation data

Somatic single nucleotide mutation data was downloaded from the Broad Institute GDAC Firehose^23^.

#### H&E deep learning lymphocyte fractions benchmark

Lymphocyte fractions were generated by Saltz et al. for 13 TCGA cancer types^14^. Multiple linear regression was applied in a similar manner as for the leukocyte methylation analysis, instead using a hyperbolic sine transformation of lymphocyte fraction as a response variable to meet normality and heteroscedasticity assumptions of the model. Models for each method were fitted using only method estimates of lymphocytes as explanatory variables. BLCA n=298, BRCA n=944, CESC n=229, COAD n=414, LUAD n=470, LUSC n=385, PAAD n=169, PRAD n=332, READ n=142, SKCM n=384, STAD n=335, UCEC n=447, UVM n=63.

#### Independent methods side-by-side benchmarks

The benchmarking validation analyses for each of the methods were replicated, where possible, to match the parameters used in the original publications. Of the six methods there were four benchmarking datasets available; either online or provided by the authors. Each of the datasets contained samples with bulk gene expression values along with matched “ground truth” values. The CIBERSORT benchmarking dataset, provided by the authors on request^5^, consisted of flow cytometry values of different immune cell types from PBMC samples. The xCell benchmarking datasets, SDY311 and SDY420, were publicly available for download from ImmPort^15^, and the validation data consisted of matching CyTOF quantification of immune cells from PBMC samples. The MCP-counter publication used gene expression profiles from GEO (accession number GSE39582) and IHC counts of CD3^+^, CD8^+^, and CD68^+^ cells (available on request from the authors)^7^. TIMER benchmark consisted of H&E stained slides from TCGA Bladder urothelial carcinoma (BLCA) study. Pathological estimations of these slides were carried out to categorise each sample into one of three categorical levels for neutrophil abundance: “Low”, “Medium” or “High”; estimations are available from the TIMER online resource^6^. For all benchmarking experiments, except TIMER, concordance was measured using correlation between “ground truth” values and the immune estimations of each method. Due to the variation in the degree of specificity to which cell subsets were defined, summations of subsets were required to allow accurate comparisons in some cases. For the TIMER benchmark, the in-silico neutrophil estimations for each method were grouped by low, medium and high pathological estimation, then compared through ANOVA with Tukey post hoc.

## DATA AVAILABILITY

All code and data generated from this study are available either on request or at https://github.com/olliecast/ConsensusTME.

## ACKNOWLEDGEMENTS

A.J.S. was supported by a doctoral fellowship from the Cancer Research UK Cambridge Institute and the Mexican National Council of Science and Technology (CONACyT). O. C. was supported by the Brown Performance Group, Innovation in Cancer Informatics Discovery Grant. M.L.M. was supported by Cancer Research UK core grant (C14303/A17197).

## SUPPLEMENTARY FIGURES

**Figure S1:**
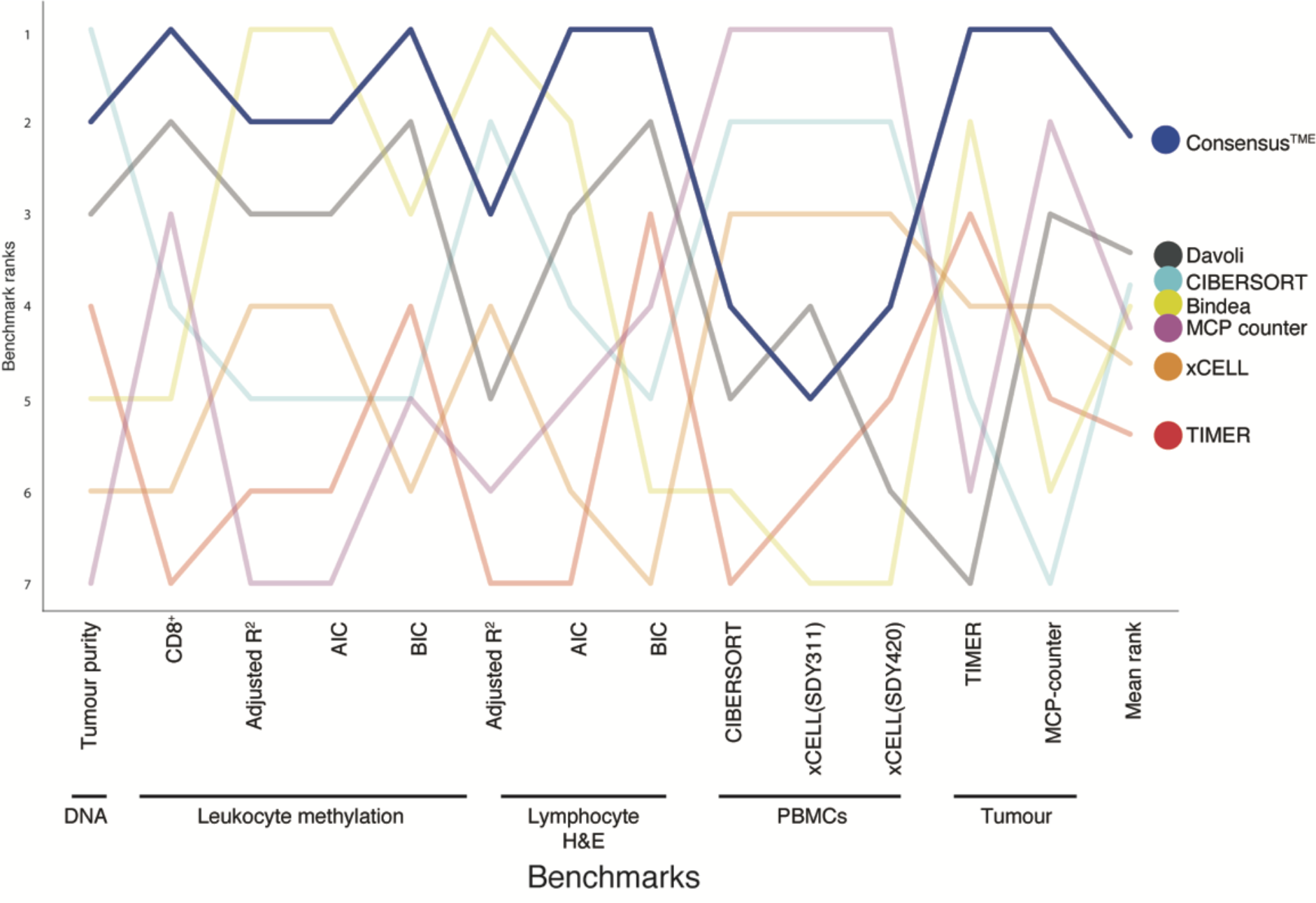
Summary of benchmarking experiments with Consensus^TME^ highlighted. Line plot for rank of method in each benchmark. Mean rank in final column. As Davoli the method does not allow for prediction of neutrophils it was given the lowest rank for the TIMER benchmark.

## CONTRIBUTIONS

*CRediT standard taxonomy* http://dictionary.casrai.org/Contributor_Roles

**Conceptualization** [Ideas; formulation or evolution of overarching research goals and aims] AJS, MM

**Data curation** [Management activities to annotate (produce metadata), scrub data and maintain research data (including software code, where necessary for interpreting the data itself) for initial use and later re-use] AJS, OC

**Formal analysis** [Application of statistical, mathematical, computational, or other formal techniques to analyse or synthesize study data] OC, AJS

**Funding acquisition** [Acquisition of the financial support for the project leading to this publication] MM

**Investigation** [Conducting a research and investigation process, specifically performing the experiments, or data/evidence collection] OC, AJS

**Methodology** [Development or design of methodology; creation of models] AJS, OC

**Project administration**[Management and coordination responsibility for the research activity planning and execution] AJS, MM

**Resources** [Provision of study materials, reagents, materials, patients, laboratory samples, animals, instrumentation, computing resources, or other analysis tools] MM

**Software** [Programming, software development; designing computer programs, implementation of the computer code and supporting algorithms; testing of existing code components] OC, AJS

**Supervision** [Oversight and leadership responsibility for the research activity planning and execution, including mentorship external to the core team] AJS, MM

**Validation** [Verification, whether as a part of the activity or separate, of the overall replication/reproducibility of results/experiments and other research outputs] OC, AJS

**Visualization** [Preparation, creation and/or presentation of the published work, specifically visualization/data presentation] OC, AJS

**Writing original draft** [Preparation, creation and/or presentation of the published work, specifically writing the initial draft] AJS, OC

**Writing review & editing** [Preparation, creation and/or presentation of the published work by those from the original research group, specifically critical review, commentary or revision] AJS, MM, OC

